# Cannabis use and risk of schizophrenia: a Mendelian randomization study

**DOI:** 10.1101/092015

**Authors:** Julien Vaucher, Brendan J. Keating, Aurélie M. Lasserre, Wei Gan, Donald M. Lyall, Joey Ward, Daniel J. Smith, Jill P. Pell, Naveed Sattar, Guillaume Paré, Michael V. Holmes

**Affiliations:** Service of Internal Medicine, University Hospital of Lausanne, Lausanne, Switzerland; Department of Surgery, University of Pennsylvania, Philadelphia, PA, United States; Centre for Psychiatric Epidemiology and Psychopathology (CEPP), University Hospital of Lausanne, Prilly, Switzerland; Oxford Centre for Diabetes, Endocrinology, and Metabolism, Churchill Hospital Campus, University of Oxford, Oxford, UK; Wellcome Trust Centre for Human Genetics, University of Oxford, Oxford, UK; Institute of Health & Wellbeing, University of Glasgow, Scotland, UK; Institute of Cardiovascular and Medical Science, University of Glasgow, Scotland, UK; Population Health Research Institute, Hamilton Health Sciences, Department of Clinical Epidemiology and Biostatistics, McMaster University, Hamilton, ON, Canada; Population Genomics Program, Chanchlani Research Centre, McMaster University, Hamilton, ON, Canada; Department of Pathology and Molecular Medicine, McMaster University, Hamilton, ON, Canada; Thrombosis and Atherosclerosis Research Institute, McMaster University, Hamilton, ON, Canada; Clinical Trial Service Unit & Epidemiological Studies Unit (CTSU), Nuffield Department of Population Health, University of Oxford, UK; Medical Research Council Population Health Research Unit at the University of Oxford, UK

**Author notes:** Corresponding authors: Dr Julien Vaucher Service of Internal Medicine Lausanne University Hospital (CHUV) 46, Rue du Bugnon 1011 Lausanne Switzerland T : 0044 77 1 777 25 24 E, Dr Michael V. Holmes Clinical Trial Service Unit & Epidemiological Studies Unit (CTSU) Nuffield Department of Population Health University of Oxford United Kingdom T : 0044 1865 74 3644 E.

## Abstract

Cannabis use is observationally associated with an increased risk of schizophrenia, however whether the relationship is causal is not known. To determine the nature of the association between cannabis use on risk of schizophrenia using Mendelian randomization (MR) analysis, we used ten genetic variants previously identified to associate with cannabis use in 32,330 individuals. Genetic variants were used in a MR analyses of the association of genetically determined cannabis on risk of schizophrenia in 34,241 cases and 45,604 controls from predominantly European descent. Estimates from MR were compared to a metaanalysis of observational studies reporting effect estimates for ever use of cannabis and risk of schizophrenia or related disorders. Genetically determined use of cannabis was associated with increased risk of schizophrenia (OR of schizophrenia for users vs. non-users of cannabis: 1.37; 95%CI, 1.09 to 1.67; P-value=0.007). The corresponding estimate from observational analysis was 1.50 (95% CI, 1.10 to 2.00; P-value for heterogeneity = 0.88). The genetic instrument did not show evidence of pleiotropy on MR-Egger (Egger test, P-value=0.292) nor on multivariable MR accounting for tobacco exposure (OR of schizophrenia for users vs. nonusers of cannabis, adjusted for ever vs. never smoker: 1.41; 95% CI, 1.09-1.83). Furthermore, the causal estimate remained robust to sensitivity analyses. These findings strongly support a causal association between genetically determined use of cannabis and risk of schizophrenia. Such robust evidence may inform public health message about the risks of cannabis use, especially regarding its potential mental health consequences.

## INTRODUCTION

Cannabis is the most widely misused illicit drug with estimated 182 million consumers in 2013 globally.^1^ Several high-profile observational studies have reported a positive, dose-dependent association between cannabis use and risk of schizophrenia, especially in young people in whom cannabis use is particularly high.^2^ The lifetime risk of schizophrenia is around 0.7%, and the natural history of disease carries a high risk of longterm symptoms and disability together with a reduced life expectancy.^3^ In addition, schizophrenia represents a high economic burden with an estimated cost of $63 billion/year in the United States.^4^ Clarifying the causal role between cannabis use and risk of schizophrenia is therefore important to understanding the health impacts of cannabis exposure and to inform on potential preventative strategies to alleviate the burden of disease from schizophrenia.^5^

A substantial body of observational evidence supports the hypothesis that cannabinoids play a role in the development of schizophrenia.^2^ Prospective observational studies, with decades of follow-up and accounting for a large number of potential confounding factors (such as demographic, family history, personal history, socio-economic or other environmental markers) have consistently demonstrated that exposure to cannabis is associated with increased risk of schizophrenia or related disorders.^2^ These findings have been reinforced by basic research experiments that point to cannabis altering various neurotransmission pathways linked to pathogenesis of psychotic disorders and by interfering with neurodevelopment in adolescents.^6^ Despite this, any causal link between cannabis use and psychotic disorders remains controversial as observational findings can always be hampered by confounding (where another risk factor associated with cannabis actually causes disease) and/or reverse causality bias (where individuals affected by schizophrenia may be more prone to consume cannabis).^2, 7^ Moreover, cannabis use is intimately associated with tobacco consumption and the latter has been observationally related to risk of schizophrenia, which may confound the link between cannabis and the disease.^8^

In the setting where a randomized trial - representing the optimal method to test a clinical hypothesis - of a harmful exposure (such as cannabis consumption) would be unethical, a genetic approach represents a valid alternative to assess causality free from confounding or reverse causality bias.^9^ Using Mendelian randomization principles, causality between an exposure (such as cannabis use) and an outcome (e.g. schizophrenia) can be tested through use of genetic markers that associate with the exposure, employed as instrumental variables providing certain assumptions are met.^10^ Recent developments of Mendelian randomization facilitate assessing the robustness of the genetic instrument by testing for presence of pleiotropy (where genetic markers associate with the outcome through more than one causal pathway, also known as horizontal, or directional, pleiotropy), Egger Mendelian randomization (MR-Egger) provides a statistical test for presence of pleiotropic effects of the genetic instrument, and a causal estimate that takes this into account, whereas multivariable MR provides a causal estimate for an exposure that statistically adjusts for a potential pleiotropic effect of the genetic marker(s) with a risk factor (e.g. tobacco consumption).^11, 12^

We used single nucleotide polymorphisms (SNPs) associated with ever use of cannabis reported in a recent genome-wide association study (GWAS),^13^ as instrumental variables to clarify the causal role of cannabis consumption on risk of schizophrenia. We then assessed for presence of pleiotropy of the genetic instrument through MR-Egger and adjusted for potential shared pathways and/or confounding by tobacco consumption in multivariable MR. We additionally conducted sensitivity analyses by restricting to SNPs with putative functional roles and by sequentially excluding each SNP from the analysis. Finally, we compared the causal estimate to a meta-analysis of observational studies.

## PARTICIPANTS AND METHODS

### Observational analysis between ever use of cannabis and risk of schizophrenia

Observational studies reporting an association between cannabis use and risk of schizophrenia were selected from a recent and comprehensive review of the literature (published in 2016) and a meta-analysis from 2007 reporting prospective studies showing an association between cannabis use and schizophrenia.^2, 14^ As only one study reported schizophrenia as an outcome,^15^ we slightly broadened our inclusion criteria to also include studies reporting related disorders (schizophreniform disorder and psychotic symptoms). To identify additional studies that may be eligible for inclusion since the meta-analysis from 2007, we conducted a PubMed search (**Figure S1**).

To compare with the causal estimate (see below), we restricted to studies that reported ever use of cannabis (compared to never users of cannabis) as an exposure and a corresponding risk estimate for schizophrenia or related disorders and identified four studies that met these criteria.^15-18^ We found one additional study in which the definition of the exposure was similar (any use of cannabis, provided that individuals have consumed cannabis ≥5 times) and also included it in the analysis.^19^ The pooled effect estimate was derived using a random-effects meta-analysis of study summary estimates. **Tables S1 and S2** summarize the main characteristics of included and excluded studies, respectively.

### Genetic markers associated with ever use of cannabis

We used the 10 leading SNPs from a recent GWAS (contributing studies outlined in **Table S3**), comprising data of participants from European ancestry predominantly, on cannabis use (phenotype defined as ever use of cannabis during participants' lifetime) to obtain the gene-exposure (SNP-cannabis) association estimates and their corresponding standard errors (SE) (**Table S4**).^13^ Although none of the SNPs surpassed a conventional genome-wide significance threshold (P-values comprised between 4.6 x 10^−7^ and 3.1 x 10^−6^ in the discovery analysis), estimates were directionally consistent across the vast majority of contributing studies (**Table S4**). These SNPs can individually, and cumulatively, be considered as valid instruments for Mendelian randomization analysis.^20^

### Association between cannabis-associated genetic markers and risk of schizophrenia

The gene-outcome (SNP-risk of schizophrenia) association estimates were obtained using the publicly available GWAS repository on schizophrenia from the Psychiatric Genomics Consortium (http://www.med.unc.edu/pgc/downloads). **Table S5** describes the contributing studies. SNPs were directly matched with the 10 SNPs associated with ever use of cannabis. The number of individuals and the relationships between datasets are presented in **Figures S2**. We used the same reference allele for each SNP to orientate cannabis and schizophrenia estimates.

### Statistical analysis

Mendelian randomization analysis was conducted by first generating an instrumental variable (IV) estimate for each SNP. The IV estimate for each SNP was generated by dividing the association of each SNP with risk of schizophrenia by the corresponding association with risk of ever use of cannabis and the standard error was estimated using the delta method.^21^ We pooled instrumental variable estimates across SNPs using a fixed-effects meta-analysis. Estimates of the association of each SNP with ever use of cannabis were not transformed. In order to generate a Mendelian randomization estimate for 'users vs. non-users' of cannabis (as opposed to a per-1-log unit increase in ever use of cannabis), we transformed the summary estimate from meta-analysis using estimates of risk of schizophrenia in the population, and the prevalence of schizophrenia in never users of cannabis, as previously described.^22^ A full description of the methodology is provided in the **supplement**.

### Characteristics of the genetic instrument

#### (i) Strength of instrument and power to detect a causal effect

In Mendelian randomization analyses, but especially in the context where multiple SNPs that did not achieve GWAS significance are used cumulatively, there are certain characteristics that need to be tested.

First, a concern might be weak instrument bias. Conventionally, when using datasets that overlap for the SNP-exposure and SNP-outcome, this can generate biased estimates and yield an inflated causal estimate (arising from correlation of the error terms of SNP-exposure and SNP-outcome).^23^ However, in our case, there was only minimal overlap (4.7%) between the datasets used to derive the effect estimates for SNPs with ever use of cannabis and risk of schizophrenia (**Figure S2**), minimizing the possibility of weak instrument bias yielding a false positive association. I.e. in the context of non-overlapping datasets, weak instrument bias result in a false negative association.^23^

We estimated instrument strength by calculating the proportion of variance in use of cannabis explained by each SNP. We then derived the F-statistic of each SNP individually and cumulatively (full details provided in the **supplement**).

We estimated power to detect the same magnitude of association reported in the observational studies, using a two-sided alpha of 0.05. Power was 100% and is presented in **Table S6**.

#### (ii) Assessment of directional pleiotropy

We tested for presence of unmeasured pleiotropy of the genetic instruments using MR-Egger as described by Bowden *et al.*^11^ Essentially, this uses the same principles of testing for small study bias in meta-analysis. The methodology was similar as for conventional Mendelian randomization analysis (described above), with the exception that all alleles (and corresponding estimates) were oriented in the direction of an increase in the exposure prior to the analyses. The standard error was obtained by bootstrap resampling 10,000 times.

### Sensitivity analyses

As tobacco consumption has been related to risk of schizophrenia and shares a strong genetic correlation with cannabis in Stringer et al,^8, 13^ we conducted a multivariable MR - to adjust for shared pathways with and/or potential confounding by tobacco - using summary statistics for the association of each of the 10 cannabis-related SNPs with tobacco (ever vs. never smokers) derived from 111,898 participants (51,984 ever smokers and 59,914 never smokers) from the UK Biobank (http://www.ukbiobank.ac.uk). Selection of participants and genotyping are described in the **supplement**. Multivariable MR was conducted by regressing the SNP-cannabis estimates on SNP-schizophrenia estimates adjusting for SNP-tobacco estimates.^12^ The standard error was obtained by bootstrap resampling 10,000 times.

We conducted two sensitivity analyses. First, we assessed the robustness of the summary causal estimate to inclusion of SNPs by sequentially removing each SNP from the Mendelian randomization analysis.

Second, we restricted the analyses to two SNPs (rs73067624 and rs4471463) located within two genes *(KCNT2* [1q31] and *NCAM1* [11q23], respectively) that were associated with ever use of cannabis in the gene-based tests of associations in Stringer *et al.*^13^ These two genes are potentially functional: *KCNT2* encodes a potassium voltage-gated channel thaï may play a role in addiction.^13, 24^ Previous studies have found that markers linked to *KCNT2* are related to cocaine dependence and opioid consumption.^24^ *NCAM1* regulates pituitary growth hormone secretion and is implicated in dopaminergic neurotransmission,^13^ and has been associated with dependence to nicotine, alcohol and heroin.^25^

All statistical analyses were conducted using Stata v.13.1 (Stata Corp, TX, USA).

## RESULTS

### Observational association between ever use of cannabis and risk of schizophrenia and related disorders

One prospective study met our primary research criteria and reported that ever use of cannabis (compared to no use) was associated with an odds ratio (OR) for schizophrenia of 1.50 (95% CI, 1.10-2.00). When meta-analysing this estimate with other prospective observational studies reporting related traits, including schizophreniform disorder and psychotic symptoms (encompassing a total of 1,326 cases and 58,263 controls), ever use of cannabis was associated with a 43% increase in the risk of schizophrenia or related disorders (OR, 1.43; 95% CI, 1.19-1.67; I^2^=0%) using random-effects modelling (**Figure 1**).

**Figure 1.**
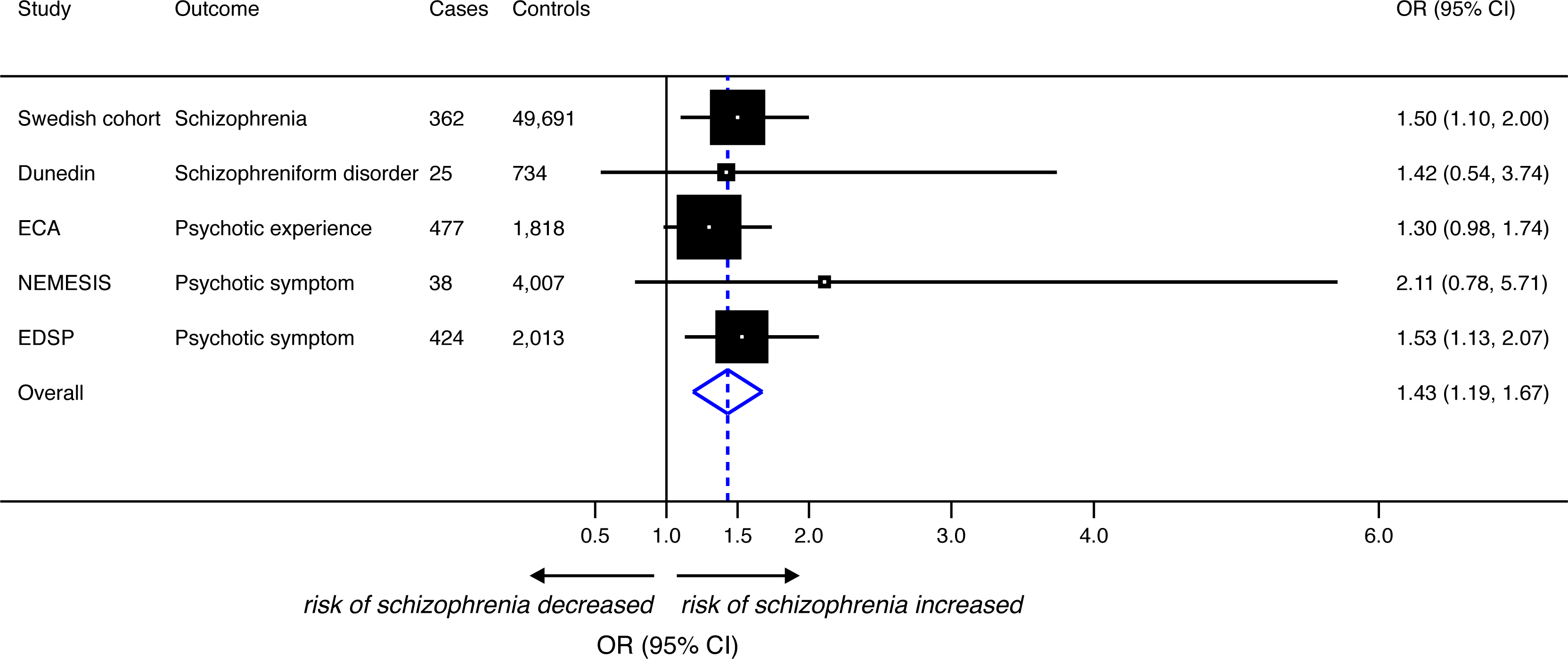
Meta-analysis of prospective observational studies reporting an association between use of cannabis and risk of schizophrenia or related disorders. Random-effects meta-analysis of observational studies. Studies are sorted by type of outcome. Odds ratios (OR) and 95% confidence intervals (CI) express the risk of schizophrenia or psychotic symptoms for ever use of cannabis (compared to never use). For additional information on each study, see **Table S1**. Dunedin, Dunedin Multidisciplinary Health & Development Study; NEMESIS, Netherlands Mental Health Survey and Incidence Study; SC, Swedish Cohort; EDSP, Early Developmental Stages of Psychopathology Study; ECA, Epidemiologic Catchment Area.

### Association between genetically determined ever use of cannabis and risk of schizophrenia

The 10 SNPs associated with ever use of cannabis explained 1.0% of its variance. There was a positive genetic association between ever use of cannabis and risk of schizophrenia (**Figure S3**). In Mendelian randomization analysis based on 34,241 cases of schizophrenia and 45,604 controls, ever use of cannabis was causally associated with risk of schizophrenia (Log OR per-1-log unit increase in ever use of cannabis, 0.07; 95% CI, 0.02-0.13; P-value=0.007) (**Figure 2**). Applying population-based estimates, this translated to a 37% increase in the risk of schizophrenia (OR for users vs. non-users of cannabis, 1.37; 95% CI, 1.09-1.67) (Figure 3).

**Figure 2.**
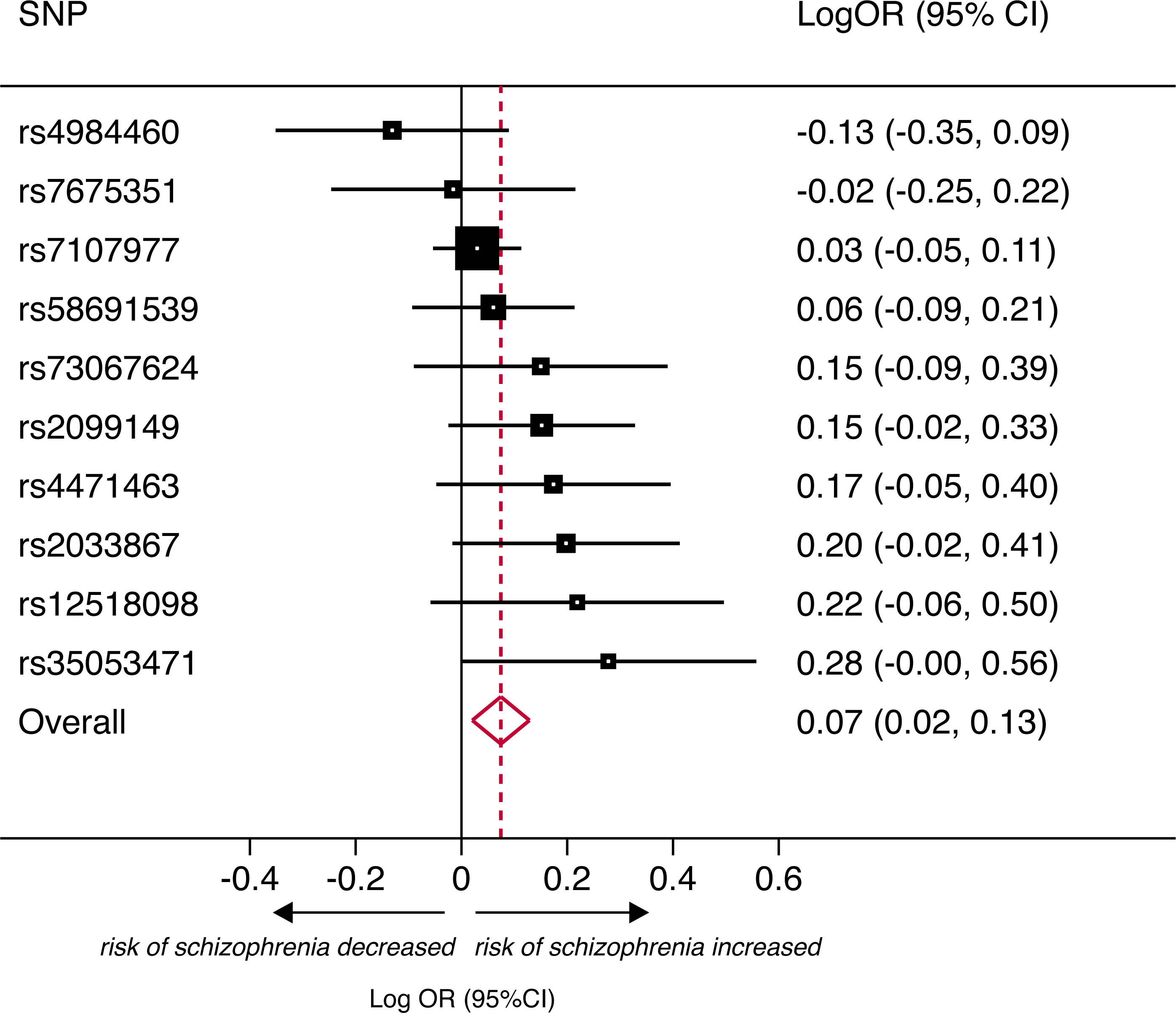
Meta-analysis of the association of genetically-determined use of cannabis and risk of schizophrenia for the 10 SNPs in the genetic instrument. Fixed-effect meta-analysis of the instrumental variable estimates for each of the 10 SNPs associated with ever use of cannabis. Log odds ratios (Log OR) and 95% confidence intervals (CI) express the risk of schizophrenia per-1-log unit increase in ever use of cannabis. The exponentiated summary estimate corresponds to an OR for schizophrenia of 1.08 (95%CI: 1.02, 1.14) per 1-log unit increase in ever use of cannabis. The method to derive the population-based OR of schizophrenia among users of cannabis compared to non-users (OR 1.55; 95%CI, 1.14, 2.00), as presented in the main text and **Figure 3**, is described in the supplement.

The Mendelian randomization estimate was consistent with estimates derived from observational analyses restricted to schizophrenia alone (test for heterogeneity, chi^2^=0.23; P-value=0.634) or schizophrenia and related disorders combined (test for heterogeneity, chi^2^=0.10; P-value=0.755) (**Figure 3**)

**Figure 3.**
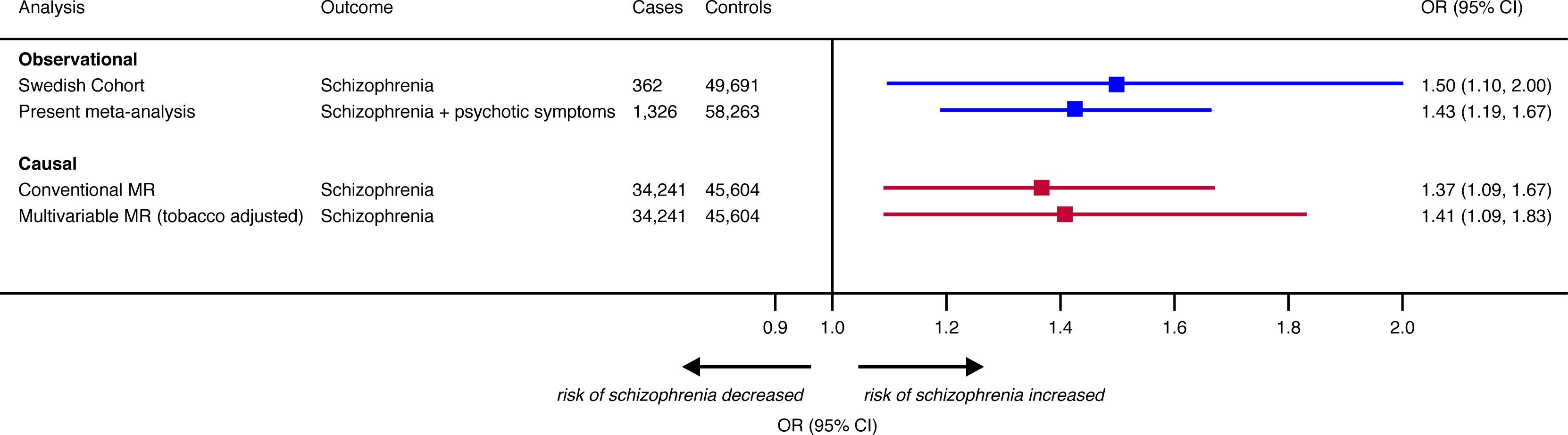
Comparison of observational (blue) and causal (red) estimates for use of cannabis and risk of schizophrenia. Two observational estimates are provided according to a stringent definition of schizophrenia (as reported in the Swedish cohort^15^) or to an outcome comprising studies reporting risk of schizophrenia or psychotic symptoms (derived from the meta-analysis reported in **Figure 1**) for ever use of cannabis. Causal estimates represent population-based associations derived by conventional (**Figure 2**) and multivariable MR. The total number of cases and controls in each analysis are presented.

### Assessment of pleiotropy of the genetic instrument

No evidence of unmeasured pleiotropy of the genetic instrument was identified using MR-Egger (P-value for pleiotropy=0.292). The estimates derived from MR-Egger and conventional Mendelian randomization are presented in **Figures S4** and **S5**.

Adjusting for the association of SNPs in the genetic instrument for smoking in multivariable MR did not show evidence of shared pathways and/or confounding with a causal estimate of schizophrenia from users of cannabis that remained stable (OR, 1.41; 95% CI, 1.09-1.83) (**Figure 3**).

### Sensitivity analyses

To further test the stability of the Mendelian randomization estimate to inclusion of SNPs that could individually distort the genetic association between cannabis use and schizophrenia, we sequentially removed each SNP from the analysis. The direction and significance of the summary association between ever use of cannabis and risk of schizophrenia remained unchanged using this approach (**Figure 4**). Furthermore, restricting the analysis to two putative functional SNPs (rs73067624 and rs4471463) showed a persistent causal association (OR for users vs. non-users of cannabis, 1.88; 95% CI, 1.00-3.21] (**Figure S6**).

**Figure 4.**
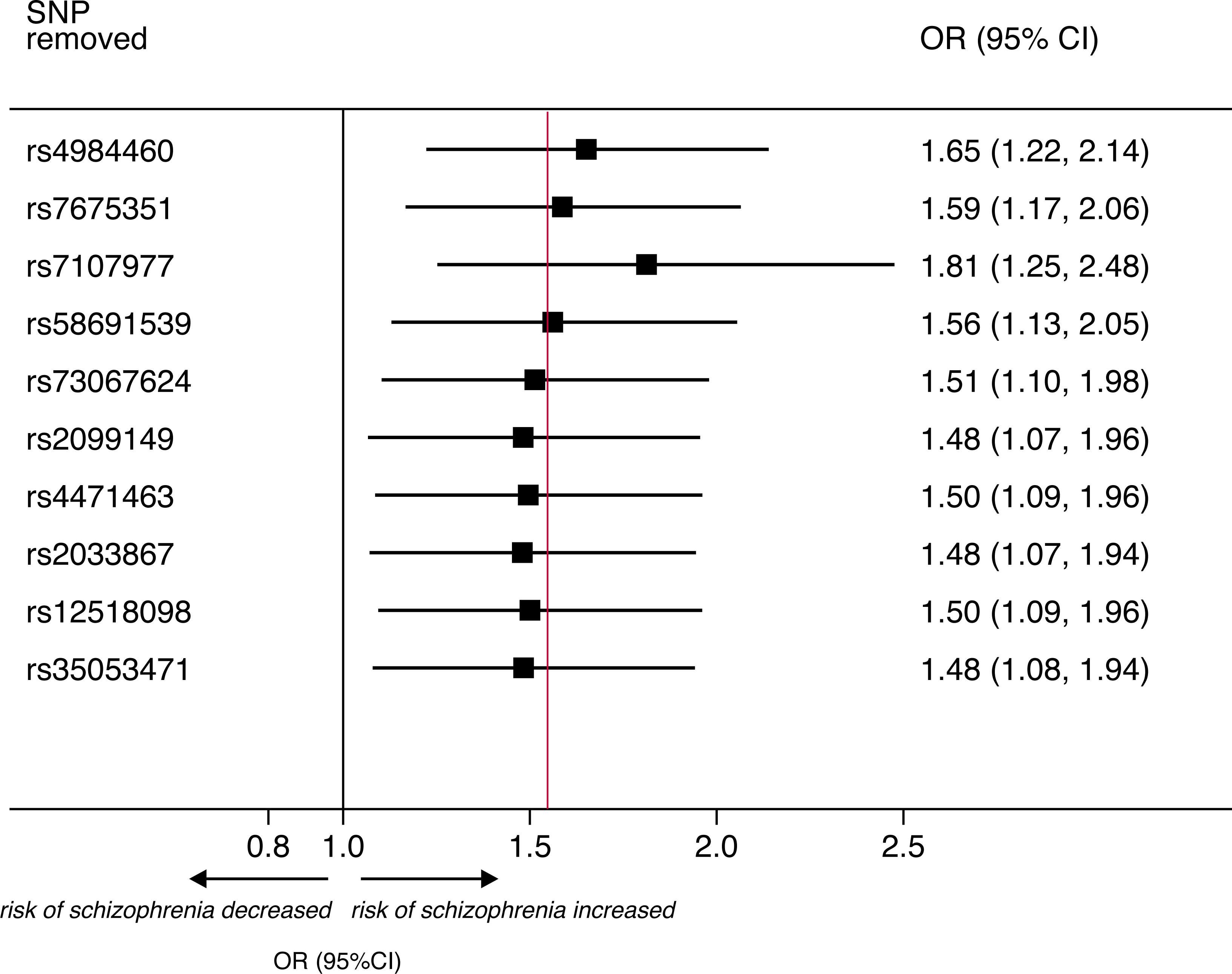
Sensitivity analysis of the association of use of cannabis and risk of schizophrenia by sequentially removing each SNP from the analysis. Plot of the Mendelian randomization summary estimates derived after sequential removal of each SNP from the analysis. The red vertical line represents the causal effect estimate (derived from Mendelian randomization) when including the 10 SNPs in the analysis (presented in **Figure 3**). Odds ratios (OR) and 95% confidence intervals (CI) represent the population-based risk of schizophrenia in users of cannabis (compared to non-users).

## DISCUSSION

This study is the first to clarify that genetically determined use of cannabis is causally associated with increased risk of schizophrenia. This finding strongly corroborates many previous prospective observational studies that identified cannabis users to be at increased risk of schizophrenia, but that could not tease out correlation from causality. As cannabis is the leading drug of misuse, this finding is timely to draw attention to the potential mental health consequences of cannabis use and to provide more robust scientific evidence to inform the public health debate on cannabis legalization.

During the last 30 years, epidemiological observations have consistently demonstrated a strong, positive and dose-dependent association between cannabis use and risk of psychotic disorders.^2, 14^ The direction and the strength of the association persisted even after adjusting for measured confounders and with long periods (~25 years) of follow-up (to attempt to minimize confounding and reverse causality bias, respectively). Our metaanalysis of prospective observational studies confirmed these findings in a magnitude that tallies remarkably closely with previous reports.^14^ Despite the consistency of observational data, clarifying whether or not cannabis use causally influences risk of schizophrenia has remained challenging. This is because observational studies, even accounting for confounding factors, can be affected by biases that can undermine the validity (such as residual confounding).^2^ As such, the ability to answer the question on causality has been at an impasse, as a randomized controlled trial, (considered the gold standard to test a hypothesis), is not possible for ethical reasons, as it would involve exposing participants to a potentially harmful exposure (a similar scenario to examining whether alcohol protects against risk of cardiovascular disease).^26^ In this setting, Mendelian randomization can provide pivotal information on causality, that can be of public health importance and inform public health guidelines.^27^ Our findings strongly support the large body of evidence from observational studies that exposure to cannabis plays a causal role in the development of schizophrenia.

Our findings are supported by studies that show that expression of schizophrenia-associated cerebral cannabinoid receptors are modified by cannabis use^28^ and that cortical maturation is altered by cannabis use in adolescents.^29^ More compellingly, small randomized trials involving human participants in laboratory conditions suggest that exposure to delta-9-tetrahydrocannabinol (THC) confers a risk to developing symptoms that mimic psychotic disorders.^6^ Observationally and genetically tobacco use is strongly correlated with cannabis use and has been proposed to act synergistically with cannabis to establish addiction.^8^ Moreover, the association between cannabis and psychotic experiences has been shown to be modulated by tobacco use, i.e. accounting for tobacco use reduces the cannabis-schizophrenia relationship.^30^ Hence, the lack of confounding by tobacco consumption, as tested by multivariable MR analysis, strengthens the findings of a primary association between cannabis use and risk of schizophrenia. Finally, our sensitivity analysis restricting to two genes with presumptive functional roles in drug dependence may suggest that cannabis affects addiction mechanisms that in turn influence the risk of schizophrenia. However, against this theory is the observation that other drugs of addiction are less associated to risk of schizophrenia or related disorders.^31^ Moreover any influence of addictive mechanisms would not undermine our findings, since cannabis exposure may be necessary to establish dependence, and addiction mechanisms could lie on the same causal pathway (**Figure S7**).

Limitations include that our study did not permit investigation of the risk of schizophrenia in relation to the quantity, type or route of administration of cannabis. Second, the precise mechanisms explaining of how some of the genetic markers under analysis alter cannabis use (or dependence) are remain unknown; however, this is not a necessary requirement to conduct a Mendelian randomization analysis using multiple loci. Third, the SNPs used in the analysis did not reach conventional genome-wide association significance thresholds. However, directions of effect were consistent in the vast majority of studies (**Table S4**) and combining individual SNPs for an analysis such as this remains valid provided the genetic instrument does not suffer from weak instrument bias. In that regard, in the context of conducting summary-level Mendelian randomization analysis using nonoverlapping data sources for the exposure and outcome (as we report here), weak instrument bias would bias the effect towards the null (i.e. opposite to weak instrument bias in overlapping datasets).^23^ This greatly increases confidence in the Mendelian randomization estimate that we report. Furthermore, our sensitivity analyses were robust to various approaches to test for stability of the causal estimates. Fourth, MR-Egger may have been underpowered to detect directional pleiotropy of the genetic instrument (if it were present).^11^ Against presence of major directional pleiotropy of any individual SNP is our analysis that excluded each SNP in turn, and to which our causal estimate remained robust. It is noteworthy that, despite these potential limitations, this study represents the closest approximation to a randomized trial on the effect of ever use of cannabis and risk of schizophrenia.

In summary, a genetic approach - representing an alternative approach to assess causality when a randomized trial would be unethical - strongly supports the notion that use of cannabis is causally related to risk of schizophrenia. This may help inform public health debate on cannabis use and preventive strategies to alleviate the burden of disease from schizophrenia.

## ACKNOWLEDGEMENT

The publicly available GWAS results on schizophrenia were obtained from the Psychiatric Genomics Consortium, University of North Carolina at Chapel Hill (https://www.med.unc.edu/pgc/results-and-downloads). The authors express their gratitude to the participants and research teams that permitted to build the publicly available dataset from the Schizophrenia Working Group of the Psychiatric Genomics Consortium, the International Cannabis Consortium, and the UK Biobank.

Sources of support: JV is supported by the Swiss National Science Foundation (P2LAP3_155086). AML is supported by the Swiss National Science Foundation (323530_151479). This research has been conducted using the UK Biobank resource which has been funded by the Wellcome Trust medical charity, Medical Research Council, Department of Health of Scottish Government, the Northwest Regional Development Agency, the Welsh Assembly Government and the British Heart Foundation.

## supplement

### Cannabis use and risk of schizophrenia: a Mendelian randomization analysis

by Julien Vaucher, MD, Brendan J. Keating, PhD, Aurélie M. Lasserre, MD, Wei Gan, PhD, Donald M. Lyall^6^, PhD, Joey Ward^6^, BSc, Daniel J. Smith^6^, MD, Jill P. Pell^6^, MD, Naveed Sattar^7^, MD, PhD, Guillaume Paré, MD, Michael V. Holmes, MD, PhD

## supplementary analysis

### The population-based risk of schizophrenia among users compared to non-users of cannabis

The method to derive the odds ratio of genetically determined use of cannabis on risk of schizophrenia in the population is based on Ross *et al.*^1^

#### Given

A. Prevalence of cannabis use in the European Union population = 0.133.^2^ This figure tallie: with prevalence of ever use of cannabis in the Swedish cohort (=0.108),^3^ which thus allows comparison between the observational and causal estimates
B. Prevalence of schizophrenia in non-users of cannabis = 0.006 (as retrieved from the Swedish cohort)^3^
C. Odds ratio for schizophrenia associated with genetically determined cannabis use (users vs. non users) at the population level
D. Genetic association with schizophrenia as function of genetic association with use of cannabis (where causal genetic effects are expressed as Log OR per allele for both schizophrenia and use of cannabis)

#### The following can be calculated

(E) Calculated population prevalence of schizophrenia: (A x B x C) + (1 - A) x B
(F) Estimated prevalence of schizophrenia in individuals with a theoretical increase in risk o use of cannabis of *e* = 2.72 fold: (A x exp(1) x B x C) + (1 - A x exp(1)) x B
(G) Estimated odds ratio for schizophrenia per *e* = 2.72 fold increase in risk of use of cannabis: F/E

#### It results that

(H) 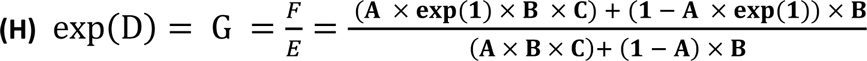 As C is the only unknown variable, the association between genetically determined use of cannabis (cannabis users vs. non-users) and risk of schizophrenia (expressed as an odds ratio) at the population level can be calculated using algebraic transformations and (H) can be simplified into:
(I) 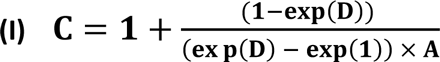

### Proportion of the variance in use of cannabis

The proportion of variance (conceptually similar to the R^2^) in use of cannabis was computed for each SNP based on the formula provided by Shim *et al.*:^4^

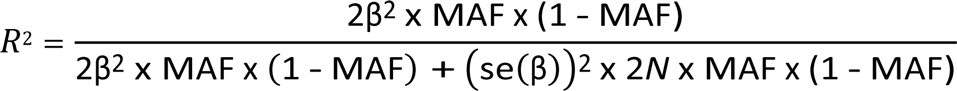

with β, effect size (beta coefficient) for a given SNP, MAF, minor allele frequency, se(ß), standard error of effect size, and *N*, sample size.

### UK Biobank: general description, genotyping, selection of participants and statistical analysis

#### General description

The UK Biobank cohort is a large prospective cohort of 502,628 participants with phenotypic information, of whom around 152,249 have genetic information available as of July 2017 with the remainder due to be released in Q3 2016.^5^ In the analysis, we reported on cross-sectional data at baseline. All participants attended one of 22 assessment centres from 2006 to 2010 where they completed a series of physical, sociodemographic, and medical assessments. Participants self-reported their smoking status as never, past or current. (A minority of n=299 refused to answer and these were removed).

#### Genotyping

UK Biobank genotyping was conducted by Affymetrix using a bespoke BiLEVE Axiom array for ~50,000 participants, and the remaining ~450,000 (for the purposes of this study 100,000) on a further updated bespoke Affymetric Axiom array (based on the 1st array). The two are extremely similar, sharing over 95% marker content. We controlled for array type as a covariate. Further information on the genotyping process is available on the UK Biobank website (http://www.ukbiobank.ac.uk/scientists-3/genetic-data), which includes detailed technical documentation (http://www.ukbiobank.ac.uk/wp-content/uploads/2014/04/UKBiobank genotyping QC documentation-web.pdf). UK Biobank provide recommendations, which we followed, for which participants to exclude from analysis based on whether: the sample failed quality control; had significant missing data or heterozygosity. We used ten (UK Biobank provided) genetic principal components to account for population stratification. All SNPs present in the current analysis were in Hardy Weinberg equilibrium.

#### Selection of participants and statistical analysis

Of the 152,249 baseline participants, there were 112,197 participants who satisfied the inclusion criteria of: passed quality control; were Caucasian; had no first cousins or closer in the cohort, and had no mismatch between reported/genetically estimated sex and ethnicity. Of these 111,898 had smoking-related data. The mean age was 56.90 (SD = 7.93) years, and 53,122 (47.4%) were male. There were 59,914 never smokers and 51,984 ever smokers (46.46%).

Risk of ever smoking per SNP (0;1;2 dose model) was adjusted for age, sex, ten genetic principal components, batch, assessment centre and array.

All analyses were conducted with PLINK and STATA v.13.

**Table S1.** Summary of the observational studies included in the analysis of cannabis use and risk of schizophrenia, presented by year of recruitment

| Study | Full name (country) | Year(s) of recruitment | Years of follow-up | Cases | Controls | Exposure (as reported in the text) | Outcome (source) | Adjustments | Reference |
| --- | --- | --- | --- | --- | --- | --- | --- | --- | --- |
| Swedish cohort | Cohort of Swedish conscripts (Sweden) | 1969-70 | 26 | 362 | 49,691 | Cannabis ever (use) | Schizophrenia (Swedish national hospital discharge) | Psychiatric diagnosis at conscription, IQ score, poor social integration, disturbed behaviour, cigarette smoking | 3 |
| Dunedin | Dunedin Multidisciplinary Health and Development Study (New Zealand) | 1972-73 | 26 | 25 | 735 | Cannabis users by age 18 | Schizophreniform disorder (DSM-IV) | - | 6 |
| ECA | Epidemiologic Catchment Area Program (US) | 1980-84 | 1 | 477 | 1,818 | Use of marijuana (vs. No use) | Self-reported psychotic experiences (Diagnostic Interview Schedule) | Sex, being in school, level of education, marital status, employment, presence at baseline of depressive episodes, manic episodes, agoraphobia and obsessive-compulsive disorder | 7 |
| EDSP | Early Developmental Stages of Psychopathology Study (Germany) | 1995 | 4 | 424 | 2,013 | Any use ($\geq 5$ times) | Psychotic symptoms (Münich version of the composite international diagnostic interview [M-CIDI]) | Age, sex, socioeconomic status, living in city, childhood trauma, predisposition to psychosis at baseline, other drug use, tobacco, alcohol, predisposition for psychosis at follow-up and depression at baseline and follow-up | 8 |
| NEMESIS | Netherlands Mental Health Survey and incidence Study (Netherlands) | 1996 | 3 | 38 | 4,007 | Baseline any use (of cannabis) | Any psychosis (based on the Brief Psychiatric Rating Scale) | Age, sex, ethnic group, level of education, unemployment and marital status, use of other drugs | 9 |

**Table S2.** Summary of the observational studies excluded from the analysis of cannabis use and risk of schizophrenia with reasons for exclusion presented by year of recruitment

| Study | Full name/description (country) | Year(s) of recruitment | Years of follow-up | Exposure (definition) | Outcome (source) | Reason(s) for exclusion | Reference (PMID) |
| --- | --- | --- | --- | --- | --- | --- | --- |
| CHDS | Christchurch Health and Development Study (New Zealand) | 1977 | 21 | Cannabis dependence (dependent vs. not dependent) based on *DSM-IV* diagnostic for cannabis dependence and frequency of cannabis use (from never use to daily use) | Psychotic symptoms (Symptom Checklist 90 [SCL-90] – 10 items) | Dependence severity (based on a count of *DSM-IV* cannabis dependence criteria) not available | Fergusson et al, Psychol Med, 2003 (12537032) and Fergusson et al. Addiction, 2005 (15733249) |
| Zürich Study | Zürich Study (Switzerland) | 1978 | 30 | Frequency of cannabis use in adolescence (3 levels : none ; casual ; regular) | Schizophrenia nuclear symptoms subscale (SCL-90-R) | Lifetime use (ever vs. never users) not available | Rössler et al, Addiction, 2012 (22151745) |
| California | California inpatient hospital admissions | 1990-2000 | 10 | Any cannabis-related ICD-9 diagnostic code within a medical recode | Readmission with any schizophrenia diagnoses (ICD-9) | Use of inpatient data - hospitalization with any record of cannabis use and subsequent hospitalization for schizophrenia. | Callaghan et al., Am J Psychiatry, 2012 (22193527) |
| ALSPAC | Avon Longitudinal Study of Parents and Children | 1991-92 | 16 | Cumulative cannabis use at age 16 years (4 levels: never; 1-20 times; 21-60 times; >60 times) | Psychotic experiences (semi-structured interview based on PLIKSi) | Lifetime use (ever vs. never users) not available | Gage et al., Psychol Med, 2014 (25066001) |
| NPMS | British National Psychiatric Morbidity Survey | 2000 | 1.5 | Cannabis use (3 levels: not used in past year, used in past year but no report of dependence; dependence) corresponding to ever use of cannabis <u>but only over the past one year</u> and cannabis dependence (dependent vs. not dependent) | Psychotic symptoms (Psychosis Screening Questionnaire) | Lifetime use (ever vs. never users) and dependence severity (based on a count of *DSM-IV* cannabis dependence criteria) not available | Wiles et al., Br J Psychiatr, 2006 (16738341) |

**Table S3.** Studies included in the cannabis-GWAS (Stringer *et al.*^10^)

| Study | Country | *N total* | Reference/PMID |
| --- | --- | --- | --- |
| *Discovery* |  |  |  |
| ALSPAC | UK | 2976 | 22507743 |
| BLTS | Australia | 721 | 23187020 |
| CADD | USA | 853 | Not published |
| EGCUT1 | Estonia | 2765 | 15133739 |
| EGCUT2 | Estonia | 970 | 15133739 |
| FinnTwin | Finland | 1029 | 23298696 |
| HUVH | Spain | 981 | 25284319 |
| MCTFR | USA | 6241 | 23363460 |
| NTR | Netherlands | 4653 | 20477721 |
| QIMR | Australia | 6778 | 17988414 |
| TRAILS | Netherlands | 1226 | 18763693 |
| Utrecht | Netherlands | 1173 | 20925969 |
| Yale Penn European American | USA | 1964 | 24166409 |
| *Replication* |  |  |  |
| Radar | Dutch | 338 | 25466800 |
| SYS | Canada | 551 | 25454417 |
| TwinsUK | UK | 2078 | twinsuk.ac.uk |
| Yale Penn African American | US | 2660 | 24166409 |

ALSPAC, Avon Longitudinal Study of Parents and Children; BLTS, Brisbane Longitudinal Twin Study; CADD, Center on Antisocial Drug Dependence; EGCUT, Estonian Genome Center University of Tartu; FinnTwin, Finnish Twin Cohort (FinnTwin12 & FinnTwin16); HUVH, Hospital Universitari Vall d'Hebron; MCTFR, Minnesota Center for Twin and Family Research; NTR, Netherlands Twin Register; QIMR, Queensland Institute of Medical Research Berghofer adults; TRAILS, TRacking Adolescents' Individual Lives Survey; Utrecht, Utrecht Cannabis Cohort (CannabisQuest); Radar, Research on Adolescent Development and Relationships ; SYS, Saguenay Youth Study.

**Table S4.**
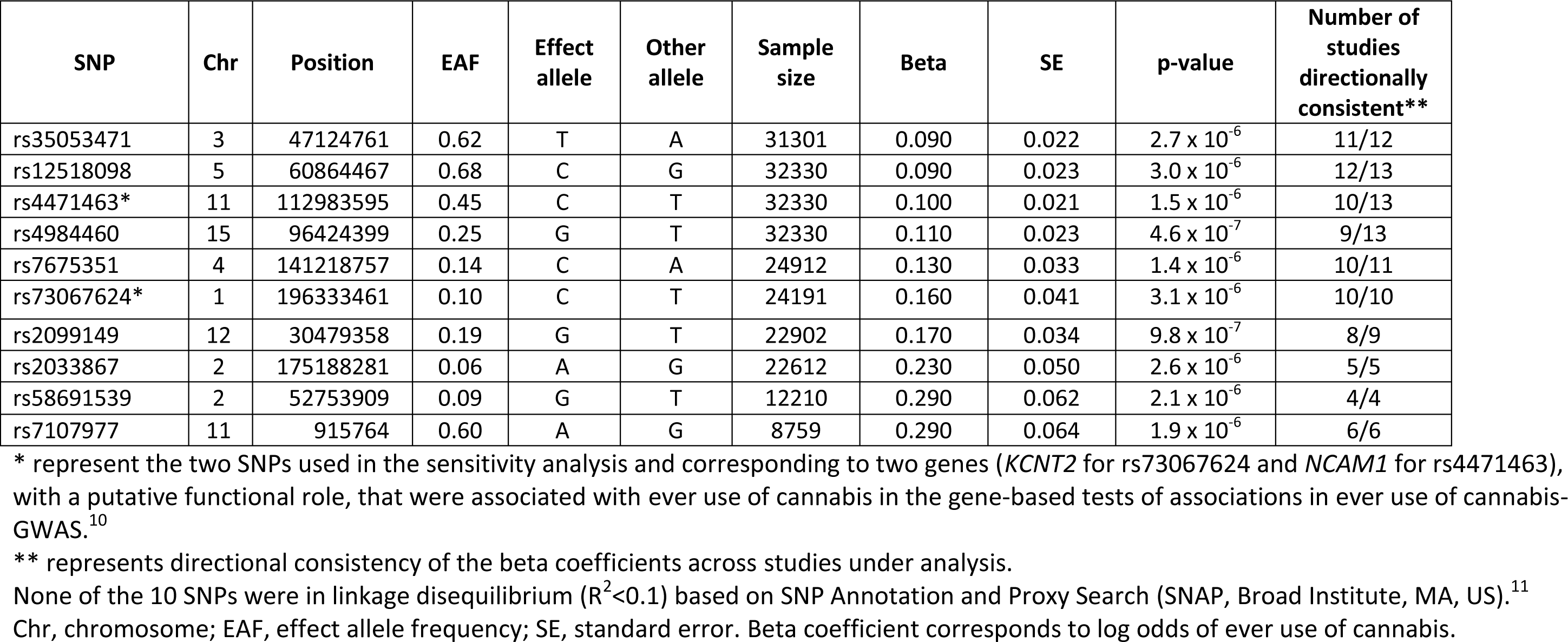
Summary of the 10 SNPs associated with use of cannabis (by increasing beta)

| SNP | Chr | Position | EAF | Effect allele | Other allele | Sample size | Beta | SE | p-value | Number of studies directionally consistent** |
| --- | --- | --- | --- | --- | --- | --- | --- | --- | --- | --- |
| rs35053471 | 3 | 47124761 | 0.62 | T | A | 31301 | 0.090 | 0.022 | $2.7 \times 10^{-6}$ | 11/12 |
| rs12518098 | 5 | 60864467 | 0.68 | C | G | 32330 | 0.090 | 0.023 | $3.0 \times 10^{-6}$ | 12/13 |
| rs4471463* | 11 | 112983595 | 0.45 | C | T | 32330 | 0.100 | 0.021 | $1.5 \times 10^{-6}$ | 10/13 |
| rs4984460 | 15 | 96424399 | 0.25 | G | T | 32330 | 0.110 | 0.023 | $4.6 \times 10^{-7}$ | 9/13 |
| rs7675351 | 4 | 141218757 | 0.14 | C | A | 24912 | 0.130 | 0.033 | $1.4 \times 10^{-6}$ | 10/11 |
| rs73067624* | 1 | 196333461 | 0.10 | C | T | 24191 | 0.160 | 0.041 | $3.1 \times 10^{-6}$ | 10/10 |
| rs2099149 | 12 | 30479358 | 0.19 | G | T | 22902 | 0.170 | 0.034 | $9.8 \times 10^{-7}$ | 8/9 |
| rs2033867 | 2 | 175188281 | 0.06 | A | G | 22612 | 0.230 | 0.050 | $2.6 \times 10^{-6}$ | 5/5 |
| rs58691539 | 2 | 52753909 | 0.09 | G | T | 12210 | 0.290 | 0.062 | $2.1 \times 10^{-6}$ | 4/4 |
| rs7107977 | 11 | 915764 | 0.60 | A | G | 8759 | 0.290 | 0.064 | $1.9 \times 10^{-6}$ | 6/6 |

\* represent the two SNPs used in the sensitivity analysis and corresponding to two genes (*KCNT2* for rs73067624 and *NCAM1* for rs4471463), with a putative functional role, that were associated with ever use of cannabis in the gene-based tests of associations in ever use of cannabis-GWAS.<sup>10</sup>
\*\* represents directional consistency of the beta coefficients across studies under analysis.
None of the 10 SNPs were in linkage disequilibrium ( $R^2 < 0.1$ ) based on SNP Annotation and Proxy Search (SNAP, Broad Institute, MA, US).<sup>11</sup> Chr, chromosome; EAF, effect allele frequency; SE, standard error. Beta coefficient corresponds to log odds of ever use of cannabis.

**Table S5.** Studies included in the Schizophrenia-GWAS (reproduced from the supplementary material of Ripke *et al.*12)

| Study | Country | Cases | Controls | Reference/PMID |
| --- | --- | --- | --- | --- |
| Umeå | Sweden | 341 | 577 | Not published |
| Umeå | Sweden | 193 | 704 | Not published |
| TOP | Norway | 377 | 403 | 19571808 |
| Uni. of Edinburgh | UK | 367 | 284 | 19571811 |
| Denmark | - | 876 | 871 | 19571808 |
| PEIC, WTCCC2 | Seven countries | 574 | 1812 | 23871474 |
| PEIC, WTCCC2 | Spain | 150 | 236 | 23871474 |
| New York, US & Israel | US & Israel | 325 | 139 | 20489179 |
| Ireland | Ireland | 264 | 839 | 19571811 |
| WTCCC2 | Ireland | 1291 | 1006 | 22883433 |
| GRAS | GRAS | 1067 | 1169 | 20819981 |
| EGCUT | Estonia | 234 | 1152 | 15133739 |
| EGCUT controls | Estonia | 347 | 310 | 15133739, 4166486 |
| EGCUT controls | Estonia | 636 | 636 | 15133739,24166486 |
| EGCUT controls | Estonia | 256 | 130 | 15133739,24166486 |
| EGCUT controls | Estonia | 1154 | 2310 | 15133739,24166486 |
| MGS | US, Australia | 2638 | 2482 | 19571809 |
| London | UK | 509 | 485 | 19571811 |
| Hubin | Sweden | 265 | 319 | 19571808 |
| Bulgaria | - | 195 | 608 | Not published |
| Toronto/Lilly (MIGen) | Canada & US | 526 | 1644 | Not published |
| Israel | - | 894 | 1594 | 24253340 |
| WTCCC controls | Six countries | 157 | 245 | 22885689 |
| New York | US | 190 | 190 | 17522711 |
| ASRB | Australia | 456 | 287 | 21034186 |
| Cardiff | UK | 396 | 284 | 19571811 |
| CLOZUK | UK | 3426 | 4085 | 22614287 |
| CLOZUK | UK | 2105 | 1975 | 22614287 |
| Netherlands | - | 700 | 607 | 19571808 |
| Finland | - | 186 | 929 | 19571808 |
| Finnish | - | 360 | 1082 | Not published |
| Portugal | - | 346 | 215 | 19571811 |
| CIDAR | US | 67 | 65 | 24424392 |
| Pfizer | - | 662 | 1172 | Not published |
| Bonn/Mannheim | Germany | 1773 | 2161 | 19571808 |
| Munich | Germany | 421 | 312 | 19571808 |
| Aberdeen | UK | 719 | 697 | 19571811 |
| CATIE | US | 397 | 203 | 18347602 |
| sw1 | Sweden | 215 | 210 | 23974872 |
| sw234 | Sweden | 1980 | 2274 | 23974872 |
| sw5 | Sweden | 1764 | 2581 | 23974872 |
| sw6 | Sweden | 975 | 1145 | 23974872 |
| CogUK | UK | 530 | 678 | 21850710 |
| NIMH CBDB | US | 133 | 269 | 11381111 |
| NIMH CBDB | US | 497 | 389 | 11381111 |
| Denmark | - | 471 | 456 | 19571808 |
| Bulgaria | - | 649 | 649 | 22083728 |
| Six countries | - | 516 | 516 | 22885689 |
| Bulgaria | - | 70 | 70 | Not published |
| Japan | - | 492 | 427 | 20832056 |
| STCRP | Singapore | 868 | 938 | Not published |
| China | - | 476 | 2018 | 24043878 |

For additional information on each study (incl. complete study name), see Ripke *et al.*<sup>12</sup> Cases included individuals with schizophrenia or schizoaffective disorder. In addition, for some studies, cases were defined based on hospital discharge records, patients having treatment-resistant schizophrenia, or registered to use clozapine.

**Table S6.** Power (two-sided α=0.05) for conventional Mendelian randomization analysis

| Exposure | Actual N<br>(Schizophrenia-GWAS) | Proportion of cases<br>(Schizophrenia-GWAS) | Observational OR | R2 of instrument | N required for 80%<br>power | Power at actual N |
| --- | --- | --- | --- | --- | --- | --- |
| Cannabis use | 79,845 | 0.429 | 1.500* | 0.010 | 18,120 | 1.0 |

Power calculation was based on the method developed by Brion et al.<sup>13</sup>
\* from Swedish cohort.<sup>3</sup>
Power at OR=1.37 (=causal estimate) is 0.99.

**Figure S1.**
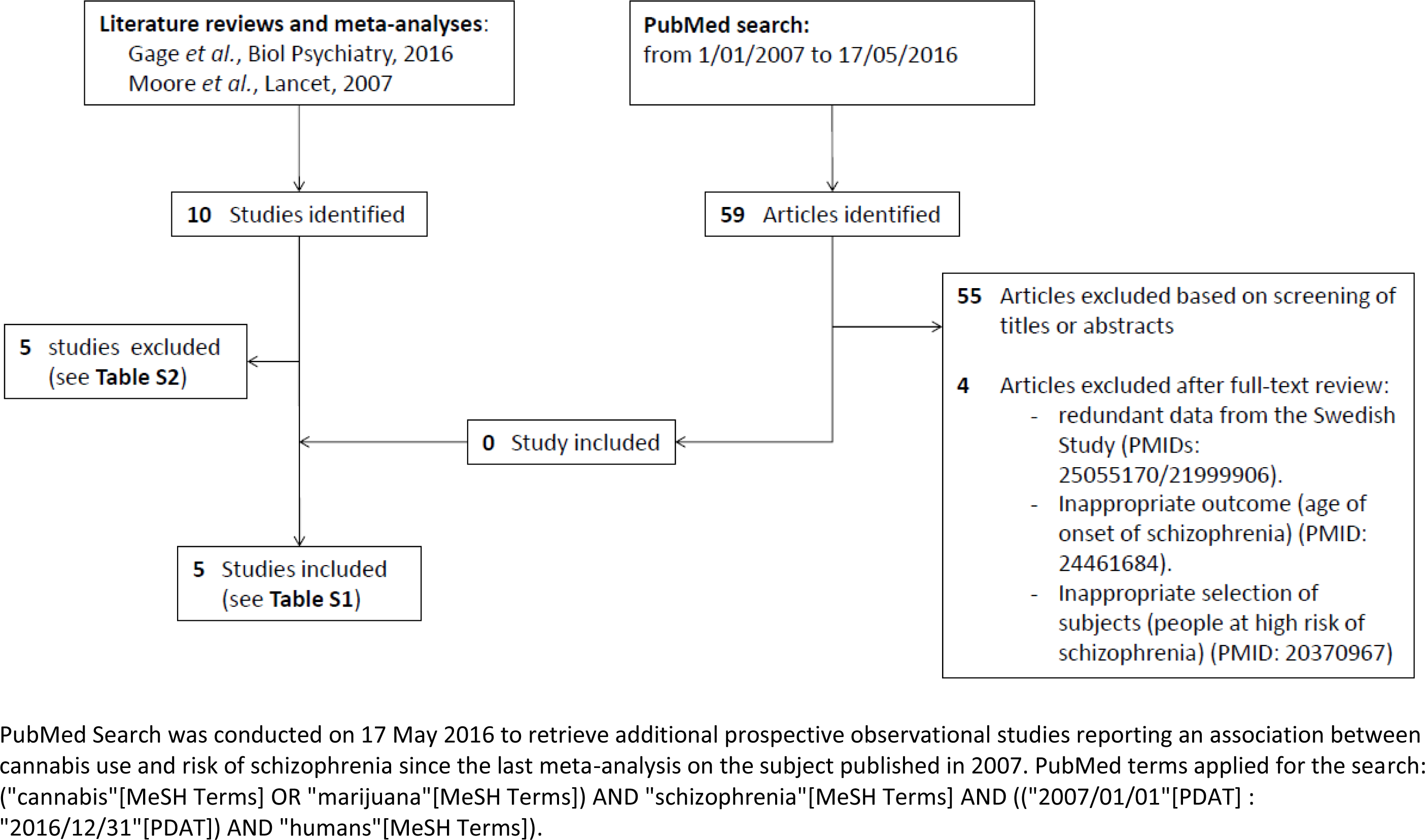
Flowchart of observational meta-analysis

**Figure S2.**
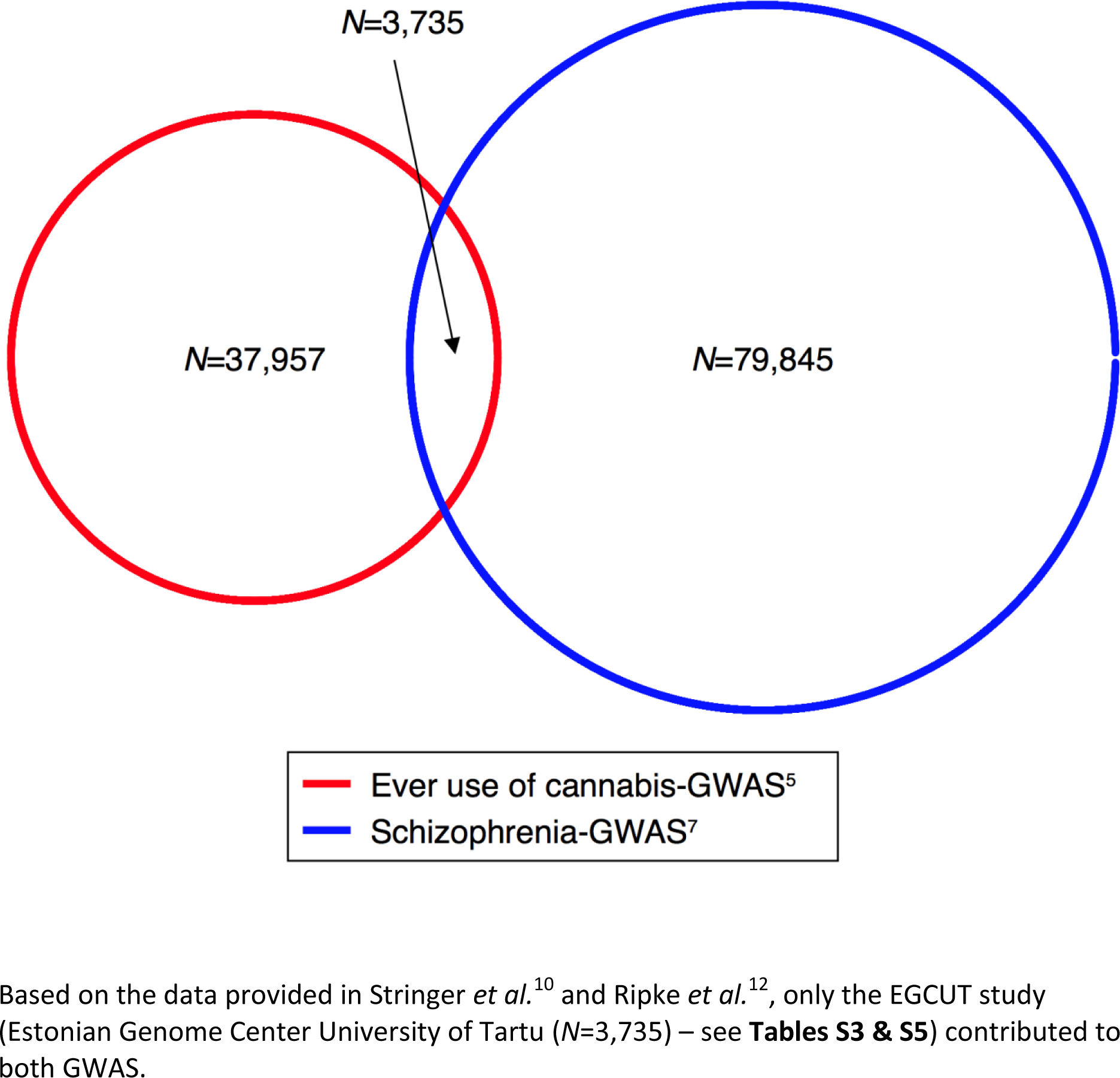
Venn diagram showing the number of individuals and the overlap between the cannabis-GWAS and the schizophrenia-GWAS

**Figure S3.**
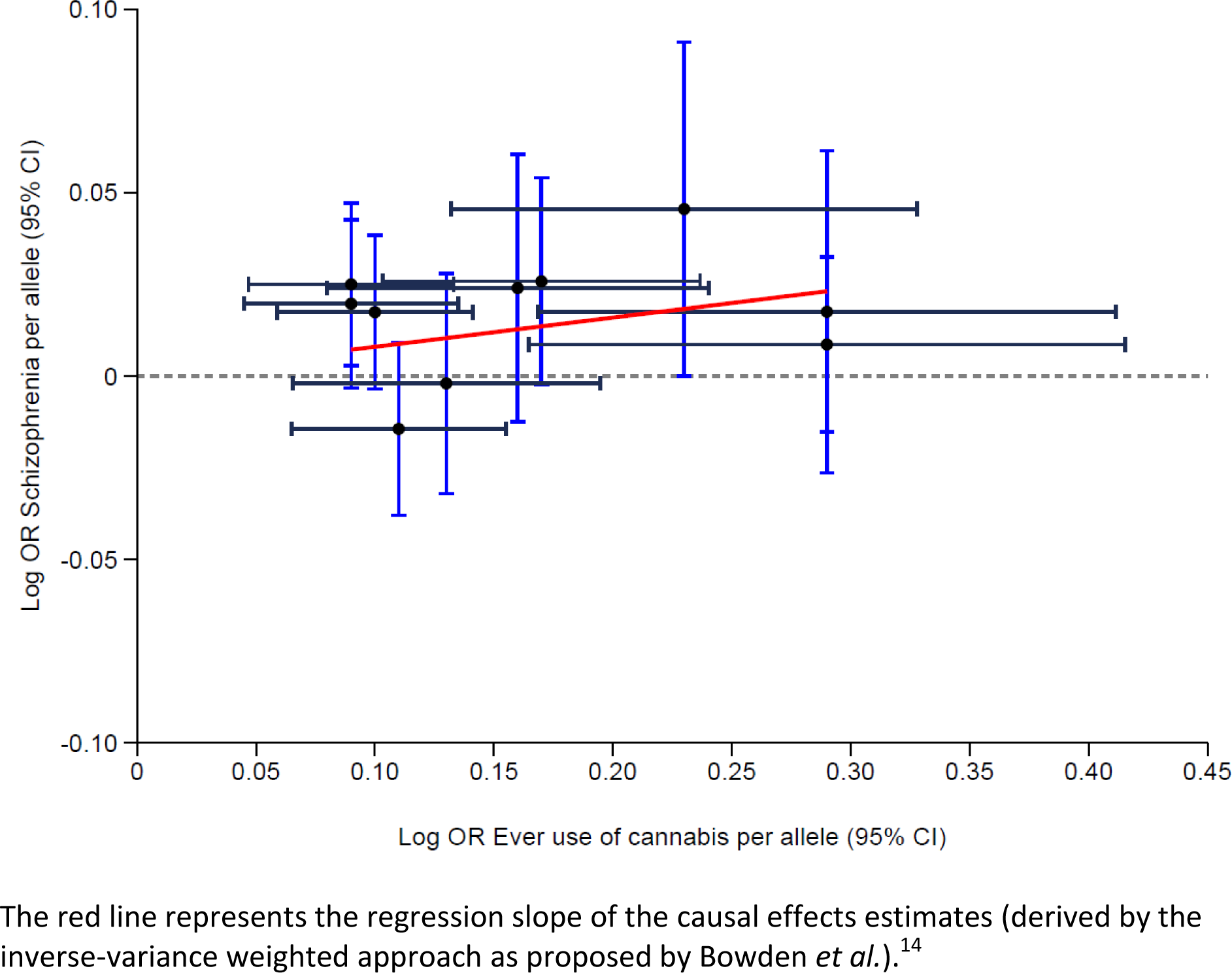
Pair-wise association plot of the 10 SNPs associated with cannabis use and risk of schizophrenia

**Figure S4.**
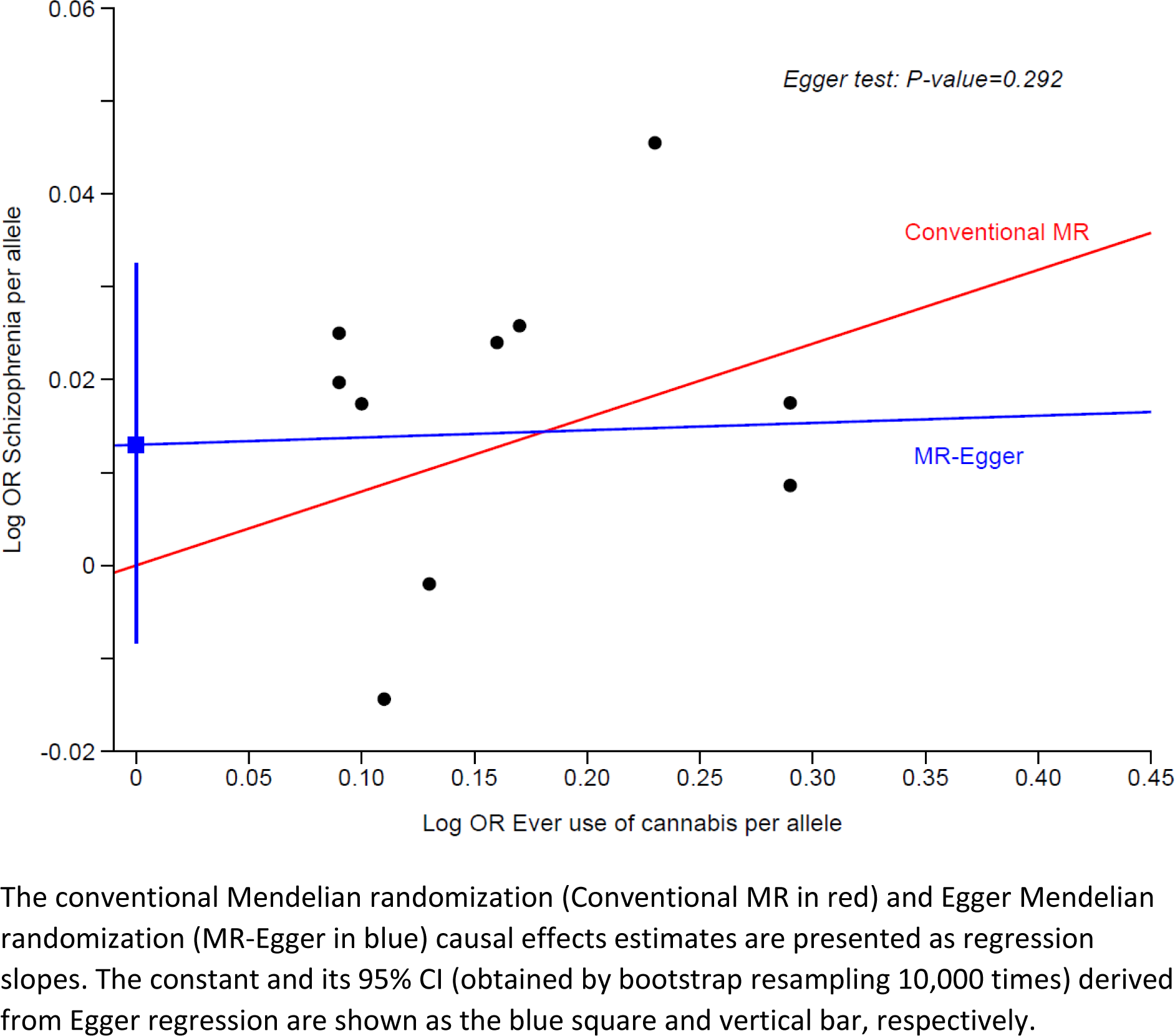
Scatter plot of the genetic association with cannabis use against genetic association with schizophrenia

**Figure S5.**
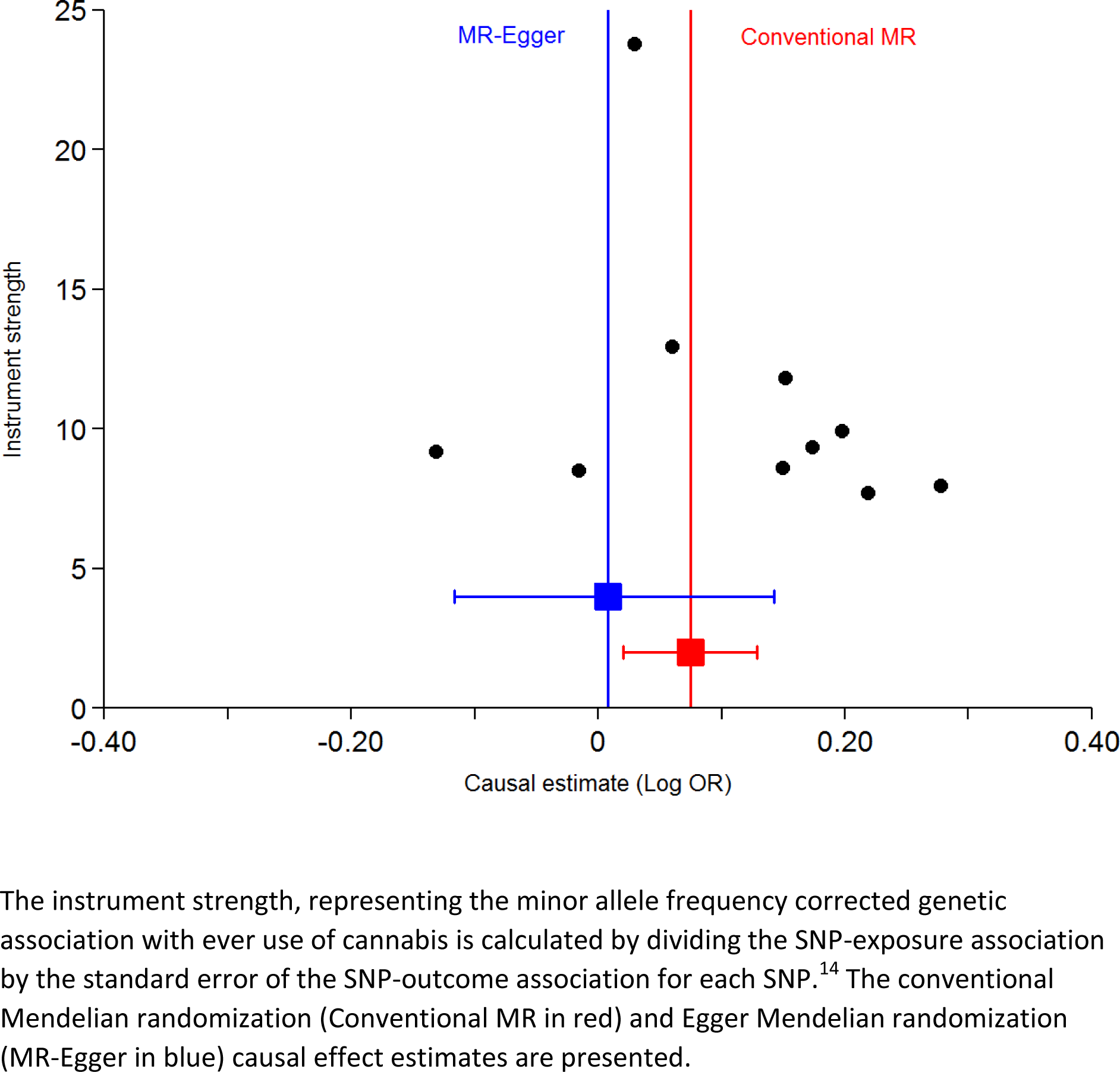
Funnel plot of the instrument strength (minor allele frequency corrected genetic association with cannabis use) against causal estimates of cannabis use on schizophrenia

**Figure S6.**
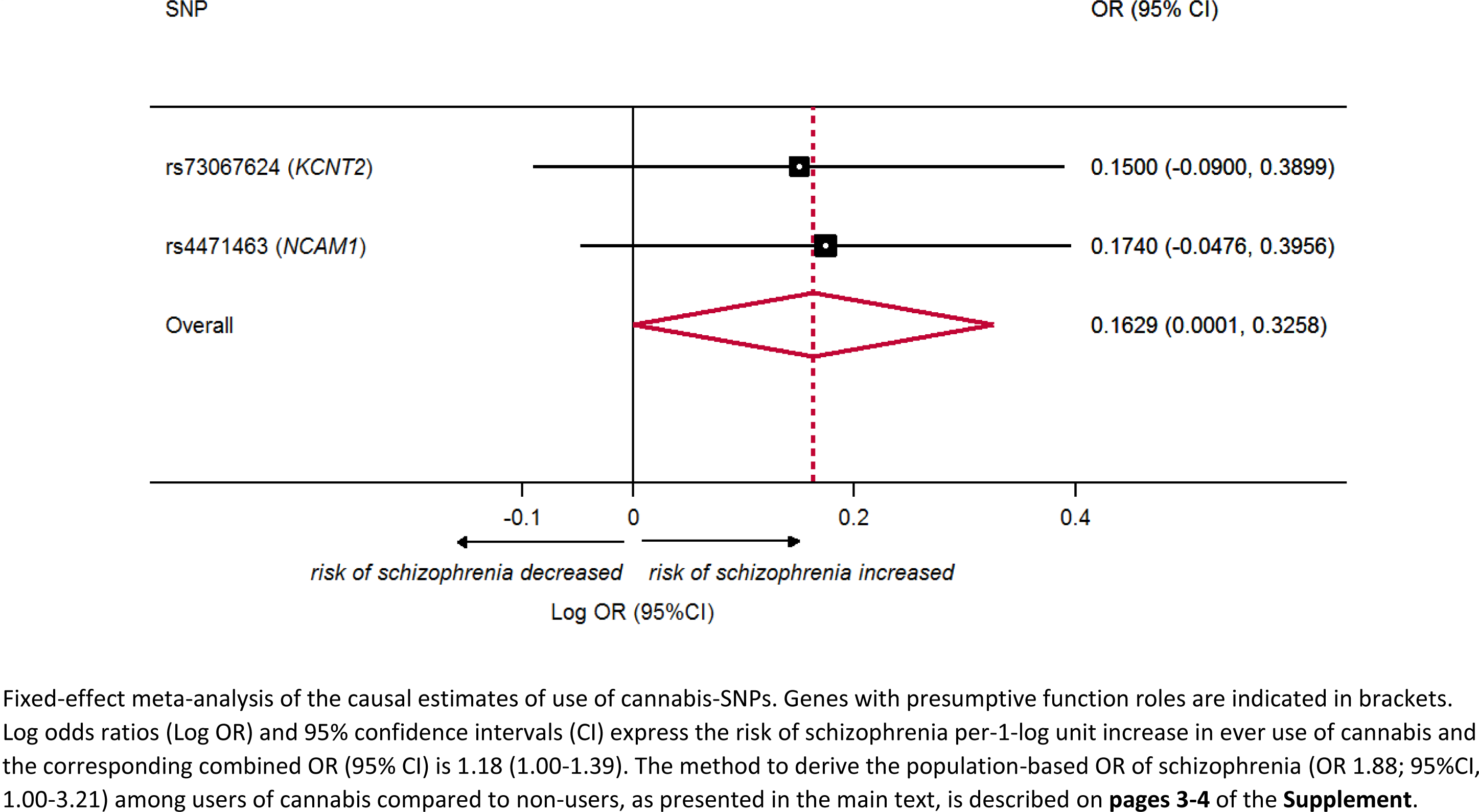
Sensitivity analyses of the association of cannabis use and risk of schizophrenia restricting to two SNPs with putative functional roles

**Figure S7.**
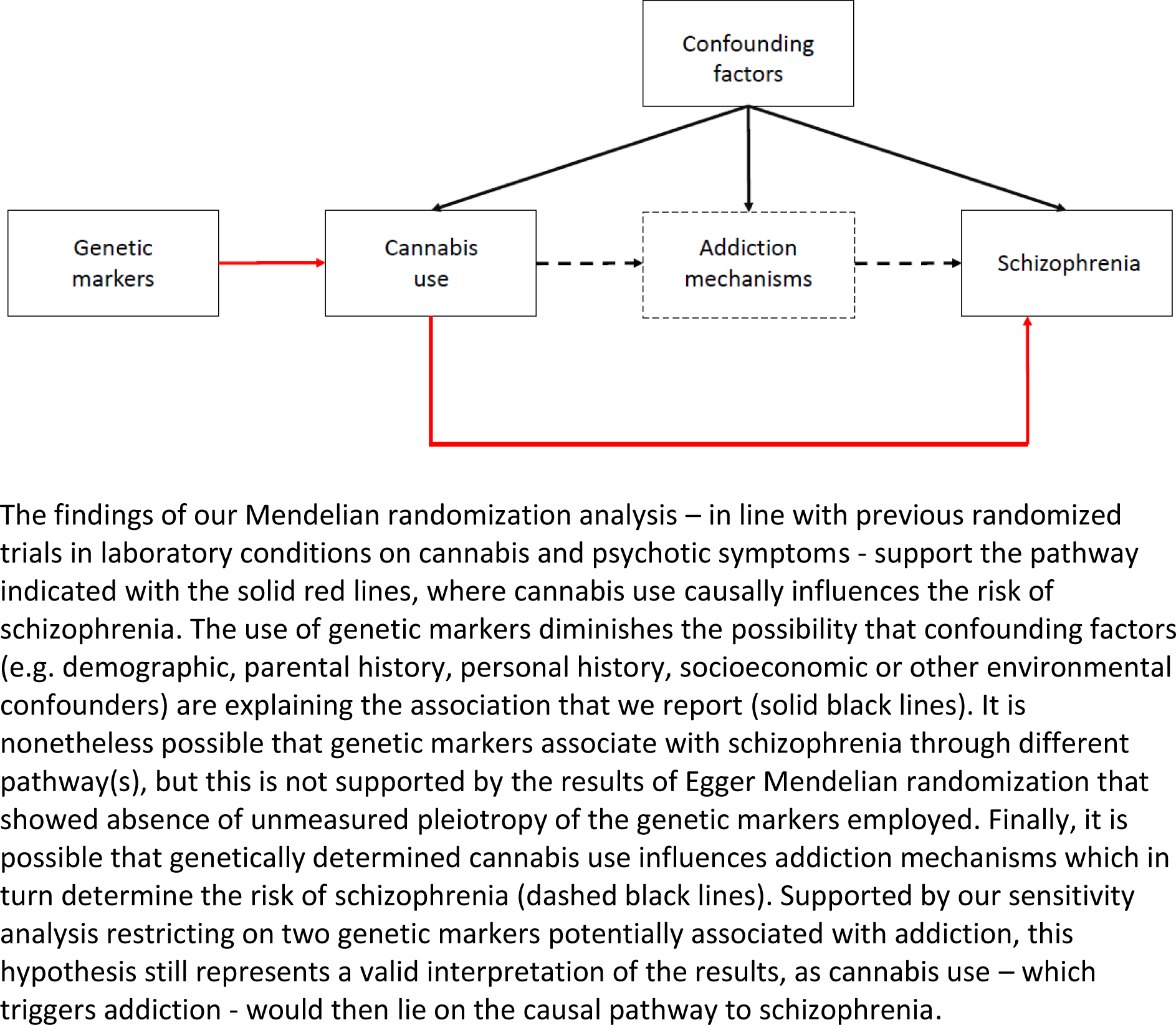
Conceptual framework representing the association between genetically determined cannabis use and risk of schizophrenia

